# Voluntary locomotion induces an early and remote hemodynamic decrease in the large cerebral veins

**DOI:** 10.1101/2024.12.02.626429

**Authors:** Beth Eyre, Kira Shaw, Sheila Francis, Clare Howarth, Jason Berwick

## Abstract

**Significance:** Behavior regulates dural and cerebral vessels, with spontaneous locomotion inducing dural vessel constriction and increasing stimulus-evoked cerebral hemodynamic responses. It is vital to investigate the function of different vascular network components, surrounding and within the brain, to better understand the role of the neurovascular unit in health and neurodegeneration.

**Aim:** We characterized locomotion-induced hemodynamic responses across vascular compartments of the whisker barrel cortex: artery, vein, parenchyma, draining and meningeal vein.

**Approach:** Using 2D-OIS, hemodynamic responses during locomotion were recorded in 9–12-month-old awake mice: wild-type, Alzheimer’s disease (AD), atherosclerosis or mixed (atherosclerosis/AD) models. Within somatosensory cortex, responses were taken from pial vessels inside the whisker barrel region ([WBR]: “whisker artery” and “whisker vein”), a large vein from the sagittal sinus adjacent to the WBR (draining vein), and meningeal vessels from the dura mater (which do not penetrate cortical tissue).

**Results:** We demonstrate that locomotion evokes an initial decrease in total hemoglobin (HbT) within the draining vein before the increase in HbT within WBR vessels. The locomotion event size influences the magnitude of the HbT increase in the pial vessels of the WBR, but not of the early HbT decrease within the draining veins. Following locomotion onset, an early HbT decrease was also observed in the overlying meningeal vessels, which unlike within the cortex did not go on to exceed baseline HbT levels during the remainder of the locomotion response. We show that locomotion-induced hemodynamic responses are altered in disease in the draining vein and whisker artery, suggesting this could be an important neurodegeneration biomarker.

**Conclusion:** This initial reduction in HbT within the draining and meningeal veins potentially serves as a ‘space saving’ mechanism, allowing for large increases in cortical HbT associated with locomotion. Given this mechanism is impacted by disease it may provide an important target for vascular-based therapeutic interventions.

## 1 Introduction

For the brain’s high energy demands to be met^1^, constant blood flow is needed. A continuous supply of blood is controlled by neurons communicating with cells of the neurovascular unit, ultimately resulting in the dilation of blood vessels across active regions of the brain^2^. This relationship between neural activity and a subsequent increase in blood flow is known as neurovascular coupling (NVC)^3,4^. Over recent years, knowledge about the pathways controlling this important mechanism has increased exponentially^5–11^. However, there are still many unanswered questions regarding the functional role of NVC across the vascular tree, which is essential for targeting pathological mechanisms along the vascular network associated with the impact of disease^12^. The brain itself is a fluid filled volume, enclosed within a hard case (the skull)^13,14^, and large intracranial pressure (ICP) changes can be ultimately devastating^15^. However, it is largely unknown how space is created within the brain to allow blood flow increases to meet metabolic demands.

Over the past decade, NVC research has transitioned from using acutely anesthetized preparations^5,6,16,17^ to performing studies in awake, behaving animals^12,18–21^. While awake imaging studies avoid the potential confounds caused by anaesthetics, these studies have their own confounds associated with behavior, including the impact of locomotion^18,22^. However, an array of behaviors can be tracked^23^, and this monitoring has allowed the field to gain a more in-depth understanding of how behaviors such as locomotion may impact NVC. For example, locomotion has been shown to induce large increases in blood flow in pial arteries and veins at its onset^20,24^, and long-lasting constrictions in dural vessels, which lie superficial to the brain surface in the outermost layer of the meninges^22^. It is thought that dural vessels likely constrict during locomotion to accommodate the locomotion-associated increased volume of arteries and veins within the brain, although it could be a myogenic response to increased blood pressure^22^. Prior work has also explored an alternative explanation of how ‘space’ is created in the brain during such large increases in cerebral blood volume (CBV), with Krieger et al.^26^ suggesting that CBV changes may be aided via the exchange of water from capillaries into neighboring tissue. However, it is worth noting that this idea is based upon modeled data and experimental support is lacking.

Despite the dynamic regulation of blood volume across the vascular tree, many vascular studies have focused on the smaller diameter cerebral blood vessels and capillaries^27,28^, and there has not yet been a systematic investigation into the effects on the large midline pial veins that drain into the sagittal sinus (referred to as draining veins)^29^. These draining veins are distinct from the cortical pial veins (e.g. here we also record from cortical “whisker” veins in the whisker barrel region of somatosensory cortex (S1)), which are part of a connected network across the cerebral microvascular bed. The anatomy of the vessels within the cerebral microvascular bed has been well characterized^30–33^, and distinct and interconnected segments have been defined (pial arterioles, penetrating arterioles, precapillary sphincters, transitional capillaries, mid-capillaries, post-capillaries and venules). Although generally assumed to be passive, remote bystanders of the vasculature, the large surface cerebral veins potentially play an important role in NVC. Early work which advocated for a passive role for veins indicated that they acted as ‘balloons’, responding only to pressure changes^34^, and more recent investigations have shown weak alpha-smooth muscle actin labeling in large cortical venules, indicating limited contractile potential^30^. However, there is some evidence that veins could in fact play a more ‘active’ role. For example, Driver et al.^35^ reported the observation of pulsatility within small cerebral veins, and veins are thought to play a role in the exchange of nutrients^36,37^, the regulation of immune cell infiltration^38^, and clearance pathways, with research suggesting that solutes empty along the venous circulation^39,40^.

To better understand how locomotion impacts hemodynamic responses across the vasculature, and whether this response is impacted by disease, we studied HbT changes (reflective of CBV) in arteries and veins from the cortical surface (within the whisker barrel region of S1), as well as veins from the sagittal sinus (draining veins) and dura mater (meningeal veins) in wild-type, Alzheimer’s disease (AD), atherosclerosis, and mixed-model (AD/atherosclerosis) mice using 2D-optical imaging spectroscopy. Initially, we collapsed the dataset across the disease groups to focus specifically on the phenomenon of the draining vein, which to our knowledge has not previously been investigated in-depth. In response to locomotion we showed an initial fast decrease in HbT in the large branches of the surface draining veins, that preceded the large increase in HbT across all vascular compartments driven by the pial arteries (“whisker artery” and “whisker vein” from inside the whisker barrel region in S1). We found that this fast draining vein HbT decrease was robust, as it was not modulated by the size of the corresponding locomotion event. We subsequently expanded our investigation of the impact of locomotion on surface vasculature to include the meningeal vessels from within the dura mater. Whilst an initial decrease in HbT was observed in both the draining and meningeal veins, interestingly, the subsequent recovery period was different between vessel types as the HbT changes did not exceed baseline levels in the meningeal vessels, but eventually showed an increase above baseline within the draining vein, likely due to the propagation of hemodynamic responses across the cortical vascular network. When we then investigated the impact of disease on the draining vein and pial vessel (whisker artery and whisker vein) responses, we showed a larger HbT increase in the artery region from the activated whisker barrel in the atherosclerosis mice versus the AD mice, and a larger HbT decrease in the draining vein in the mixed model (AD/atherosclerosis) group compared to the AD mice. Therefore, the initial draining vein response represents a previously unreported, fast, remote neurovascular signal that may be important in the overall regulation of NVC, especially to large bilateral increases in brain metabolism, which may be indicative of venous dilatory capacity and thus may be important to study in the context of neurodegeneration.

## 2 Methods

### 2.1 Animals

118 imaging sessions taken from 40 male and female mice aged between 9-12m were included in the study. Four groups of mice were used: the APP/PS1 (B6.C3-Tg (APPswe, PSEN1dE9)85Dbo/ Mmjax #34829)^41^ Alzheimer’s model, wild-type (WT) littermates, an atherosclerosis model (male only): WT-littermates injected with rAAV8-mPCSK9-D377Y (6 × 10^12^ virus molecules/ml) (Vector Core, Chapel Hill, NC) at 11 weeks of age (i/v or i/p + a western diet from 12 weeks (21% fat, 0.15% cholesterol, 0.03% cholate, 0.296% sodium; #829100, Special Diet Services UK)), and a mixed disease group (male only): APP/PS1 mice injected with rAAV8-mPCSK9-D377Y (6 × 10^12^ virus molecules/ml) (Vector Core, Chapel Hill, NC) at 11 weeks of age (i/v or i/p + a western diet from 12 weeks). Prior to surgery mice were housed in groups where possible, and were singly housed if there was no available littermate.

After surgery all mice were individually housed to ensure the whiskers and head-plate with thinned skull remained intact for imaging. Food and water were available ad-libitum (western diet was restricted to 5g per day) and mice were housed on a 12hr dark/light cycle (lights on 06:00-18:00). All experiments were carried out during the ‘lights-on’ hours. All procedures were approved by the UK Home Office and in agreement with the guidelines and scientific regulations of the Animals (Scientific Procedures) Act 1986 with additional approval received from the University of Sheffield licensing committee and ethical review board. The following study is reported in accordance with the ARRIVE guidelines. The experimenter was blinded to genotype (where possible) during experiments, and blinded during analysis. No a priori sample size calculations were conducted.

### 2.2 Surgery

Animals were anesthetized with ketamine (50mg/kg) and medetomidine (0.65mg/kg) (s/c) with isoflurane used to maintain surgical plane of anesthesia (0.5-0.8% in 100% oxygen). Carprofen (10mg/kg) was administered prior to a scalpel being used to remove hair from the head. Animals were positioned in a stereotaxic frame (Kopf Instruments) and eyes protected with Viscotears (Novartis). Body temperature was monitored and maintained at 37 degrees C with the use of a rectal thermometer and a homeothermic blanket (Harvard Apparatus). Iodine and bupivacaine (50-100mcL at 0.025%) was applied prior to revealing the skull. Suture lines were covered with cyanoacrylate glue and a dental scraper was used to score the contralateral side of the skull, to increase headplate stability. A dental drill was used to thin the bone over the right somatosensory cortex (∼4mm^2^). The skull was thinned until the pial vasculature was observed (Figure 1b). Cyanoacrylate glue was spread across the thinned region. A metal headplate was attached using dental cement (Superbond C & B; Sun Medical). Atipamezole (2mg/kg in 0.3ml warm sterile saline s/c) was given at the end of the procedure to reverse the effects of medetomidine. Following surgery, mice were placed in an incubator (29 degrees C) and monitored. Post-surgery, mice were housed individually and given at least one week to recover prior to habituation and imaging. Mice were closely monitored for weight loss and signs of pain for 3 days post-surgery and given carprofen (10mg/kg) in jelly for at least 1-day post-surgery.

**Fig. 1.**
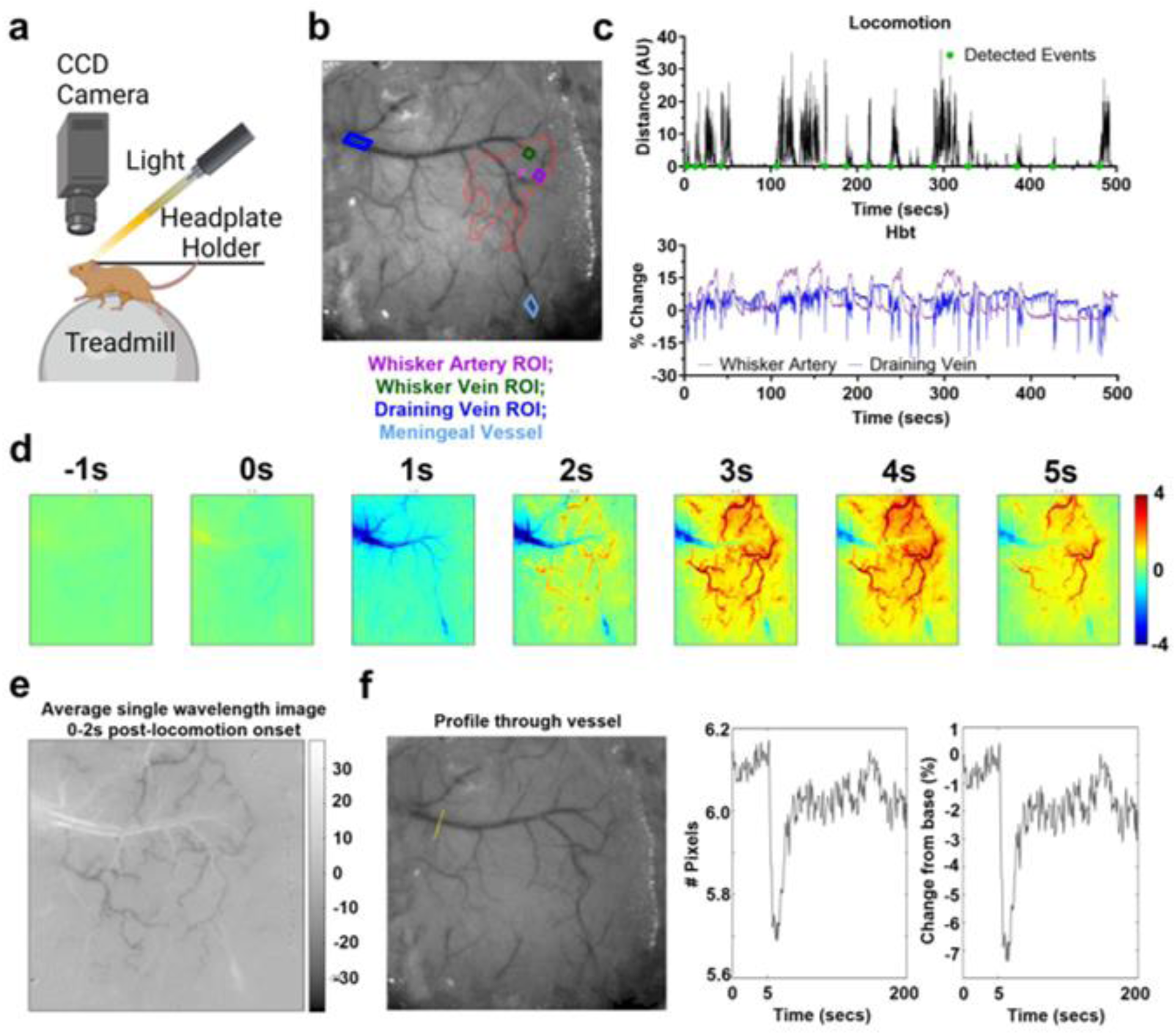
Experimental Set-up. **A.** Schematic of the 2D-OIS imaging set-up for recording of spontaneous hemodynamic responses in an awake mouse on a spherical treadmill with locomotion tracked (Created in BioRender. Eyre, B. (2025), BioRender publication link to be provided before publication). **B.** Thinned cranial window from a representative animal with cortical surface vasculature visible (imaged at 575nm illumination). The whisker barrel region is indicated by the red ROI. The purple ROI indicates the artery, the green ROI the vein, and the pink ROI the parenchyma selected from within the whisker barrel region, the dark blue ROI represents the draining vein adjacent to the whisker barrels, and the teal ROI the meningeal vessel from the overlying dura mater. **C.** Example continuous time series traces for locomotion (black, top) and HbT from the whisker artery (purple, bottom) and draining vein (blue, bottom) from one single 750s recording (first 500s visualized). The green dots on the locomotion plot represent the detected locomotion events. **D.** Spatial map generated from this representative imaging session for trial-average changes in concentration (uM) of total hemoglobin (HbT) in response to locomotion (from 1s before to 5s after onset). The scale bar represents the uM change for activated pixels, with a decrease in HbT shown in blue and an increase in red. The draining vein is observed on the left side of the images, with the rapid decrease in HbT (blue) seen in this vessel from 1s, whereas the whisker barrel arterial network is highlighted in red as HbT increases in these vessels from 2s onwards. **E.** In our illustrative example, the image of the thinned cranial window from one single wavelength captured under 2D-OIS was averaged for the 2s following detected locomotion events. The white borders along the edges of the draining and meningeal veins indicate vasoconstriction. **F.** A profile of the vessel diameter was also taken for the 2s following the onset of locomotion across the yellow line indicated in the left image, with the draining vein having darker pixels compared to the surrounding tissue. The change in the number of dark pixels representing the size of the draining vein is visualized as number of pixels (center) or a percentage change from the baseline size (right), indicating the draining vein is constricting following locomotion onset.

### 2.3 2-Dimensional Optical Imaging Spectroscopy (2D-OIS) in Awake Mice

One week after surgery mice were habituated to the awake imaging set up via increasing exposure to the experimenter and experimental apparatus (see ^42^; Figure 1a). Mice were briefly anesthetized with isoflurane (3-4%) to get them onto the awake imaging apparatus. All mice underwent between 1 and 4 2D-optical imaging spectroscopy (2D-OIS) sessions lasting approximately 30 minutes to 1 hour, which included the collection of 750 second spontaneous experiments where mice were head-fixed on top of a spherical treadmill with an optical motion sensor attached so that the impact of locomotion on hemodynamic responses could be assessed.

Widefield imaging was used to investigate blood volume changes across the surface vasculature (in somatosensory cortex and overlying dura matter). 2D-OIS uses light to assess changes in oxygenated (HbO), deoxygenated (HbR) and total hemoglobin (HbT) in the surface vessels of the cortex. To measure cortical hemodynamics, four different wavelengths of light (494 ± 20 nm, 560 ± 5 nm, 575 ± 14 nm and 595 ± 5 nm) were shone on to the thinned window, using a Lambda DG-4 high-speed galvanometer (Sutter Instrument Company, USA). A Dalsa 1M60 CCD camera was used to capture remitted light at 184 × 184 pixels, at a 32 Hz frame rate, this provided a resolution of ∼ 75 µm.

2D-OIS can also be used to create 2D spatial maps of micromolar changes in HbO, HbR and HbT, revealing the vasculature of the cortex. This is achieved by using the path length scale algorithm (PLSA) to complete a spectral analysis^43,44^. The PLSA works by using the modified Beer Lambert Law, with a path-length correction factor, as well as predicted absorption value of HbT, HbO and HbR. The relative concentration estimates were acquired from baseline values, where the concentration of hemoglobin within tissue was estimated as 100 µM, and tissue saturation of oxygen estimated at 80% in the whisker region, 90% in the whisker artery (art), 70% in the whisker vein (wv), 80% in the parenchyma, 70% in the draining vein (dv) and 70% in the meningeal vein (mn).

### 2.4 Regions of Interest (ROI) from 2D spatial maps

Custom, in-house MATLAB scripts were used to create ROIs from 2D spatial maps generated using 2D-OIS. A whisker region was generated by selecting the region of the cortex with the largest change in HbT to a 5Hz 2s whisker stimulation^42^ directed towards the left side whiskers (due to the thinned window being on the right side of the skull) in experiments conducted prior to the spontaneous locomotion recordings. This allowed for identification of the whisker barrel region within the somatosensory cortex. The code assessed pixels as being ‘active’ if their value was >1.5 STD across the whole of the spatial map. Therefore, the whisker ROI represented the area of the cortex with the largest HbT response to a 2s whisker stimulation. A ‘whisker artery’ (a pial artery inside the already defined whisker region) and ‘whisker vein’ (a pial vein inside the already defined whisker region) were then manually selected from inside the whisker ROI based on the identification of vessels from their morphology. The above was completed for each imaging session, with care taken to select the same whisker artery and vein across days for each individual mouse. In order to generate the draining vein ROI, the spatial images taken during a representative locomotion event were loaded into our in-house software within MATLAB, and the draining vein ROI manually selected based on a principal component analysis (PCA) to identify the pixels showing the greatest changes during locomotion. The draining vein was a large cerebral vein, protruding from the midline which was outside of the ‘whisker region’. The meningeal vessels were chosen from a subset of the experiments, again using PCA analysis, which often revealed the vessel in the same PCA as the draining vein in the midline of the cranial window.

### 2.5 Data Analysis

For the spontaneous recording taken during each imaging session, a continuous trace was generated for the locomotion data, and hemodynamic (HbO, HbR, HbT) responses across the whisker artery, whisker vein, and draining vein (Figure 1b-c). For each recording spatial maps were generated (see representative example in Figure 1d), where the PCA analysis revealed distinct responses in the draining veins (emerging from the sagittal sinus on the left side of the image), and the whisker barrel region (indicated by the red pixels) following locomotion onset. Where visible, a continuous hemodynamic trace was also extracted for the meningeal vein (57 sessions from 21 mice). A timed trigger point was generated for both the locomotion (recorded using optical motion sensor spherical treadmill) and hemodynamic (recorded using 2D-OIS equipment) traces in order to temporally align the corresponding traces. The locomotion trace was plotted in arbitrary units which displayed stationary periods as zeros, and movements as increasingly larger integers relating to the speed of the treadmill. Locomotion events (n=2343 total) were manually detected from the continuous time series plots, and the onset of each locomotion event marked and indexed using an in-house MATLAB graphical user interface (GUI). Both locomotion and hemodynamic traces were cut around the 5s preceding and 20s following the onset of the locomotion event. All locomotion and hemodynamic traces were normalized to their own baseline period using the following equation:

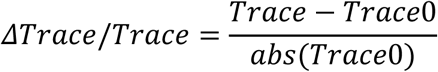

where Trace represents the entire locomotion or hemodynamic time series (i.e. from the 5s before to the 20s after the locomotion onset), and Trace0 represents the baseline period (for the 5s before locomotion onset only). For hemodynamic data, responses were then converted to a % change from baseline by multiplying by 100.

For each detected locomotion event and corresponding hemodynamic trace, the corresponding animal ID, imaging session ID, vessel type (whisker artery, whisker vein, draining vein, meningeal vein), and disease group label were extracted to allow for between group comparisons. A custom-built in-house MATLAB function was used to extract the time series metrics across groups (e.g. vessel types, locomotion conditions, or disease model) including the area under the curve, maximum peak, time to maximum peak, minima, and time to minima – which were assessed for the 0-5s after locomotion onset. The area under the curve was calculated using the ‘trapz’ function in MATLAB, and the maximum peak or minima by detecting the largest positive or negative value and calculating the average for the 0.25 seconds either side of this value (to reduce the impact of ‘noise spikes’). The time to peak was the time taken (in seconds) to reach this maximum (or minimum) value. As larger peak values would impact this timing calculation (as differences in timing could just reflect larger maximum peak or minima values), a normalized speed metric was calculated by dividing the size of the peak (maximum or minima (absolute values taken to avoid confounds of integers below zero), % HbT change) by the time to reach this value (seconds), meaning a larger positive value reflects a larger HbT value being divided by a smaller time value. Where groups were separated by the size of the locomotion event (i.e. Figure 3, top 10% vs bottom 10% locomotion trials), the area under the curve values for each locomotion event were sorted in ascending order using the MATLAB function ‘sort’.

In the representative recording session used to visualize the constriction of the draining vein (Figure 1), the constriction was calculated by averaging the spatial image for the 2s after all detected locomotion events and detecting if the number of pixels overlying the draining vein (darker pixels) changed from baseline (i.e. a reduction in the number of dark pixels indicates a constriction of the vessel has occurred).

### 2.6 Statistical Tests

Data were collapsed across disease groups for the first 4 figures, where the primary aim was to investigate the previously unreported draining vein phenomenon, and data were separated by disease group for Figure 5 to explore the impact of disease (AD, atherosclerosis, mixed AD/atherosclerosis) on this response. Statistical tests were conducted in RStudio and GraphPad Prism (Version 10), and figures created in GraphPad Prism and MATLAB (R2024a). The threshold for statistical significance was set at p ≤ 0.05. All data are presented as mean ± SEM or individual data points unless otherwise stated. The number of animals, imaging sessions and locomotion events are reported across all comparisons. Detailed statistical reports are included in the supplementary information (tables SR1-SR7).

Two-group comparisons (Figure 2f, Figure 4) were assessed for normality using a Shapiro-Wilk or Kolmogorov-Smirnov test and variance using an F-test (in line with ARRIVE guidelines), before non-parametric Wilcoxon-Signed Rank tests were conducted for paired samples (Figure 2) and Mann-Whitney tests for unpaired samples (Figure 4). Whereas multi-group comparisons (across vessel type in Figure 2, locomotion group and vessel type in Figure 3, and disease group in Figure 5) were conducted using linear mixed models, with animal ID inputted as the random factor to account for variations between groups being driven by a single outlier animal. Where post-hoc comparisons were conducted to explore significant effects, the Tukey method was used with correction for multiple comparisons. For correlational analysis exploring the relationship between the size of locomotion and hemodynamic responses across the vessel types (Figure 3a-b) a Pearson’s R correlation was conducted to assess whether the variables were significantly linearly related, and to compare the linear relationship between vessel groups (Figure 4c) a linear regression was conducted to compare the slope and intercepts.

**Fig. 2.**
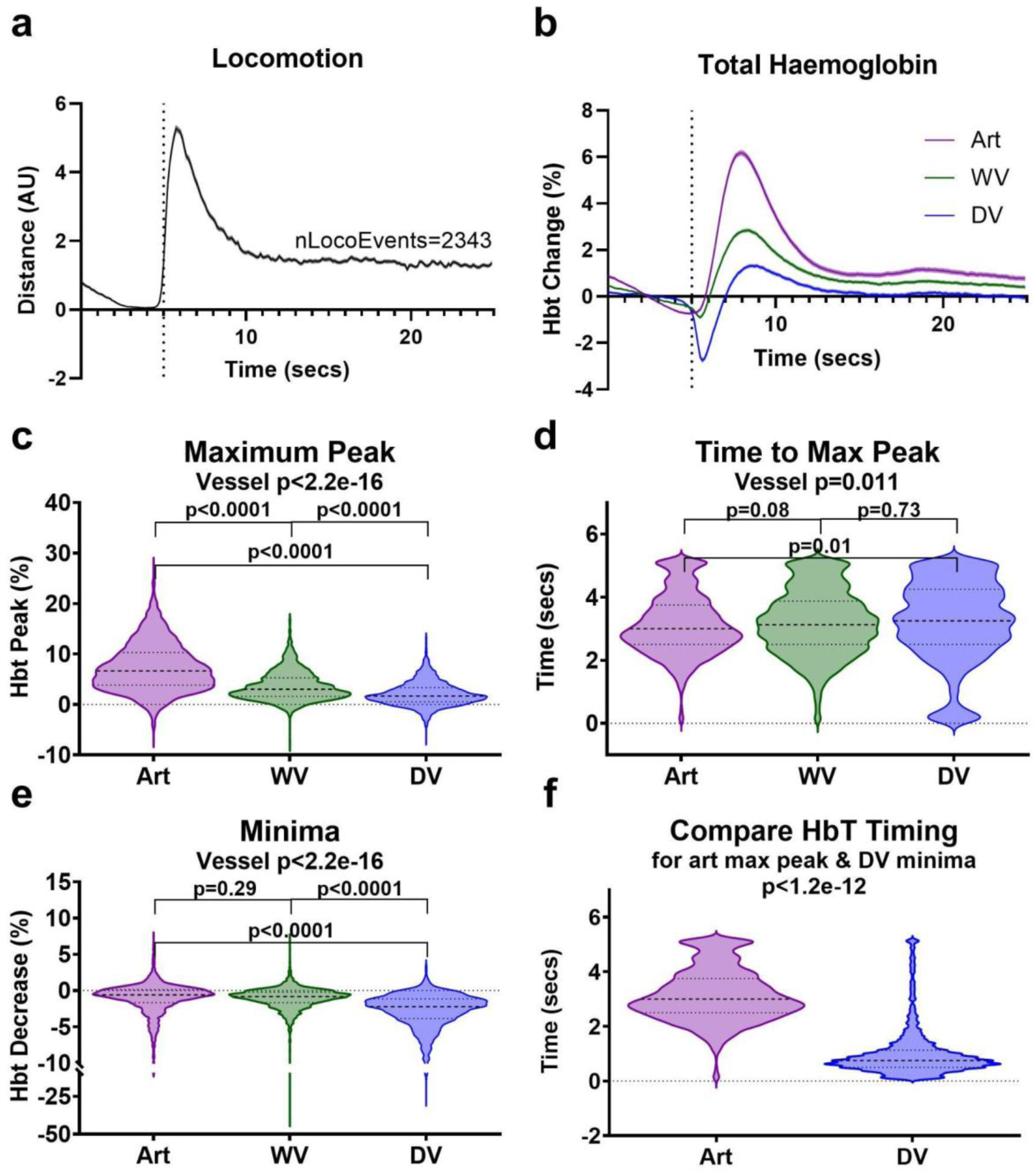
Spontaneous locomotion induces a fast, remote decrease in the large cerebral draining vein that precedes the regional HbT influx. **A.** 2343 individual locomotion events were detected across 118 imaging sessions taken from 40 mice. The 5s prior to the locomotion event, and 20s after the onset were taken, and the locomotion traces averaged. **B.** Concurrent total hemoglobin (HbT) traces corresponding to these locomotion events were extracted for the artery (purple), whisker vein (green), and draining vein (blue) in somatosensory cortex. The size and timing of the maximum peak was significantly impacted by vessel type, with the artery showing **C.** the largest increase in HbT from baseline following the onset of locomotion (art M: 7.33, SD: 4.77, wv M: 3.61, SD: 2.89, dv M: 2.06, SD: 2.40; art – dv, p<0.0001; art – wv, p<0.0001), and **D.** the quickest time to reach its maximum peak (art M: 3.13, SD: 0.96, wv M: 3.19, SD: 1.03, dv M: 3.22, SD: 1.27; art – dv, p=0.01; art – wv, p=0.08). **E.** The draining vein showed a significantly larger minimum decrease below baseline in HbT (art M: -0.97, SD: 2.01, wv M: -1.06, SD: 1.73, dv M: -2.77, SD: 2.56; art – dv, p<0.0001; wv – dv, p<0.0001), **F.** an active response which occurred significantly faster than the artery HbT increase (p<1.2e-12). P-values are taken from linear mixed-effects models with vessel type inputted as fixed-effect factor, and animal ID as the random effect, except for f) where a Wilcoxon-Signed ranks was conducted for a two-group paired comparison (see Statistics Report Table SR1). Shaded error bars represent mean +/-SEM. Horizontal lines on violin plots show median and interquartile range. Source data are provided as a Source Data file.

**Fig. 3.**
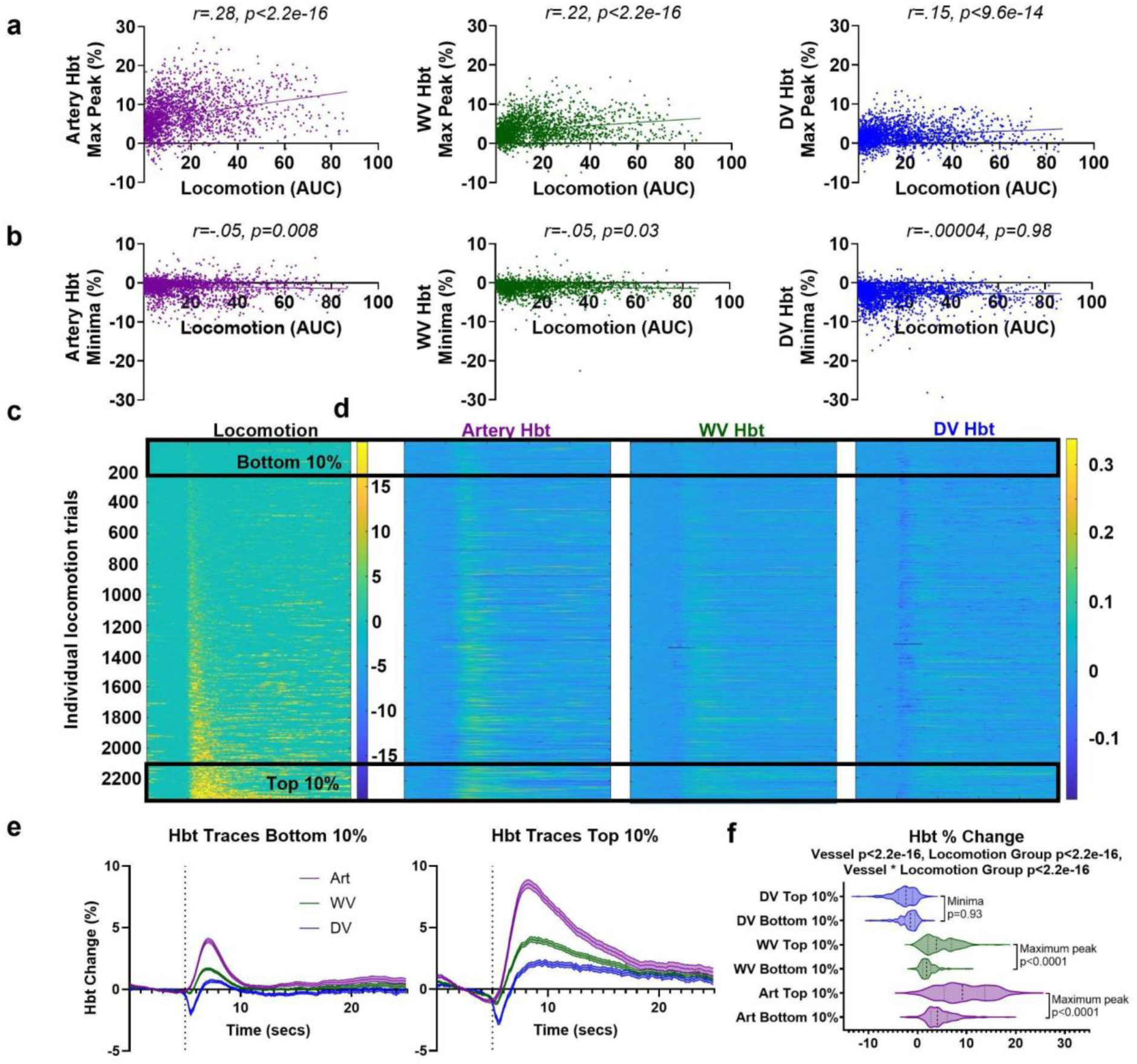
The amount of locomotion modulates the size of the HbT increase, but not the initial HbT decrease in the draining vein. **A.** The size of the locomotion event, measured using the area under the curve for the 5s following locomotion onset, correlated with the magnitude of the increase in HbT across all vessel regions (artery purple, whisker vein (WV) green, draining vein (DV) blue). **B.** The size of the initial decrease in the draining vein (measured by the minima) was not modulated by the size of the locomotion event. For correlation analysis, p-values are taken from a Pearson’s correlation (cor.test package RStudio). **C.** All locomotion events (n=2343) were sorted in ascending order from the smallest area under the curve value during the 5s following locomotion onset (5-10s), **D.** and this index was used to order the corresponding HbT traces for the artery (left), whisker vein (middle), and draining vein (right). **E.** The average HbT traces were visualized across the vessel types for the bottom (n=234) and top (n=234) 10% of locomotion trials, **F.** and whilst a significant impact of locomotion group (bottom vs top) was observed on the size of the peak across vessel types (maximum for art & wv, minimum for dv), there was no impact of locomotion group on the size of the minimum decrease in the draining vein (bottom 10% dv min peak - top 10% dv min peak p=0.93). P-values are taken from linear mixed-effects models with vessel type and locomotion group inputted as fixed-effect factors, and animal ID as the random effect (lmer package RStudio), and pairwise comparisons (with correction for multiple comparisons) were conducted using the Tukey method (emmeans package RStudio) (see Statistics Report Table SR2-4). Shaded error bars represent mean +/- SEM. Horizontal lines on violin plots show median and interquartile range. Source data are provided as a Source Data file.

**Fig. 4.**
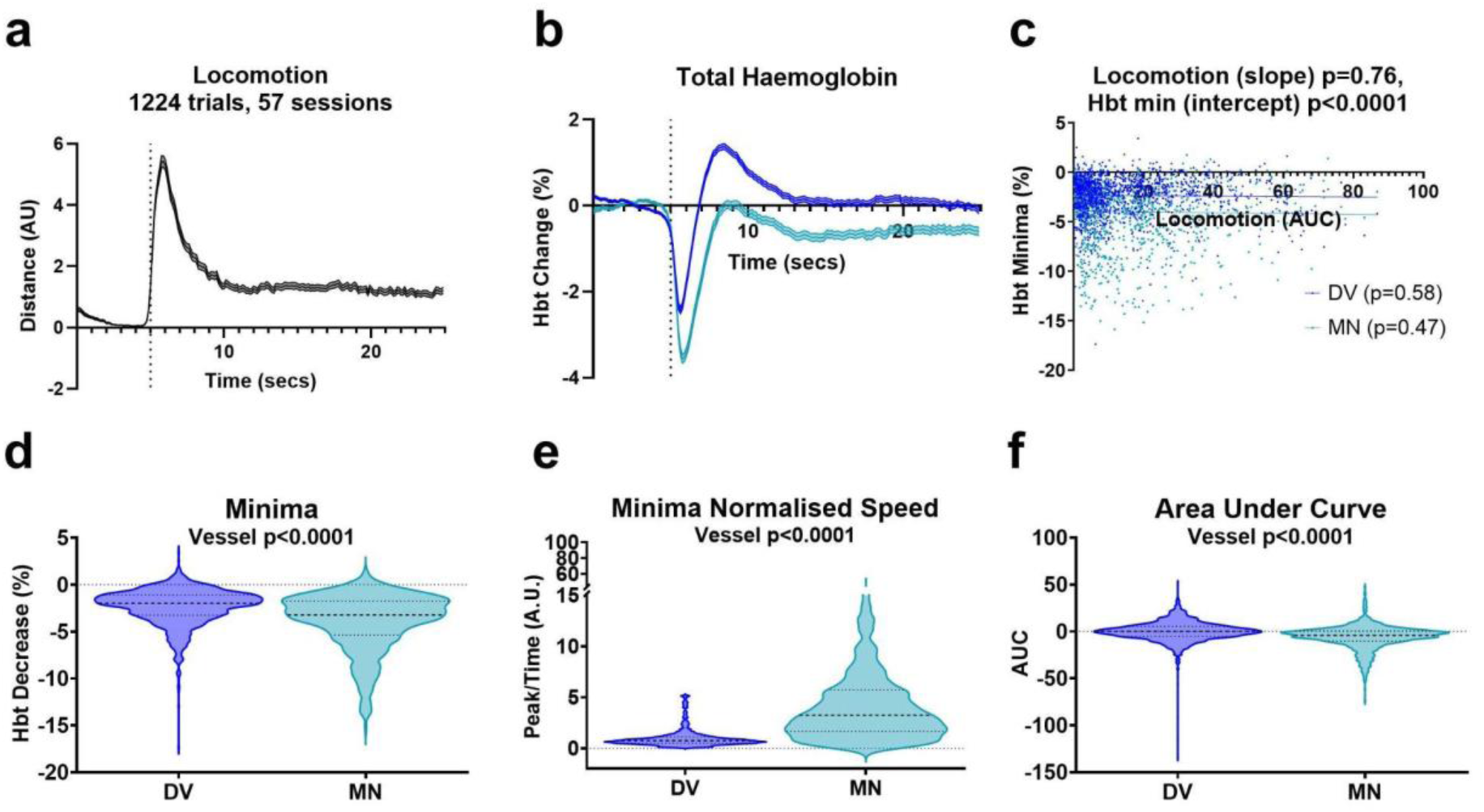
The meningeal vein shows a larger HbT decrease following locomotion onset, which unlike in the draining vein does not subsequently increase above baseline. **A.** The average locomotion traces for the 1224 locomotion events detected from the 57 imaging sessions (21 mice) where both the meningeal and draining vein were visible in the imaging window. **B.** Locomotion-dependent corresponding average total hemoglobin traces were plotted for the draining and meningeal veins, which both showed a large decrease in HbT immediately following the onset of locomotion, but with the meningeal vessel (which lies outside of the brain) not showing the subsequent overall increase in HbT during the remainder of the locomotion event. **C.** A linear regression analysis revealed there was no significant differences in the relationship between the size of the locomotion event and the size of the HbT minima (test of difference between slopes p=0.76), with neither vessel type showing a significant correlation between the size of these responses (dv p=0.58, mn p=0.47). **D.** The size of the decrease in the minima in the meningeal vessel is significantly larger than in the draining vein (dv M: -2.77, SD: 2.56; mn M: -3.97, SD: 3.06), **E.** and when we calculated a normalized speed metric by dividing the absolute value of the HbT minima (%) by the time to reach this minima (seconds), this was significantly larger for the meningeal vein which reflects a larger HbT peak being reached in a relatively faster time. **F.** In order to capture the vascular dynamics regarding the return to baseline, the area under the curve in the 5 seconds following locomotion onset was assessed between draining and meningeal veins (dv M: -0.03, SD: 10.77; mn M: -6.11, SD: 12.73), with a significant difference found as the meningeal vessel was primarily negative (sustained decrease not exceeding baseline) and the draining vein closer to 0 (as the negative decrease was followed by a subsequent increase in HbT). Two-group unpaired comparisons were conducted using Mann-Whitney tests, and linear relationships were estimated using a linear regression (see Statistics Report Table SR5-6). Shaded error bars represent mean +/- SEM. Horizontal lines on violin plots show median and interquartile range. Source data are provided as a Source Data file.

**Fig. 5.**
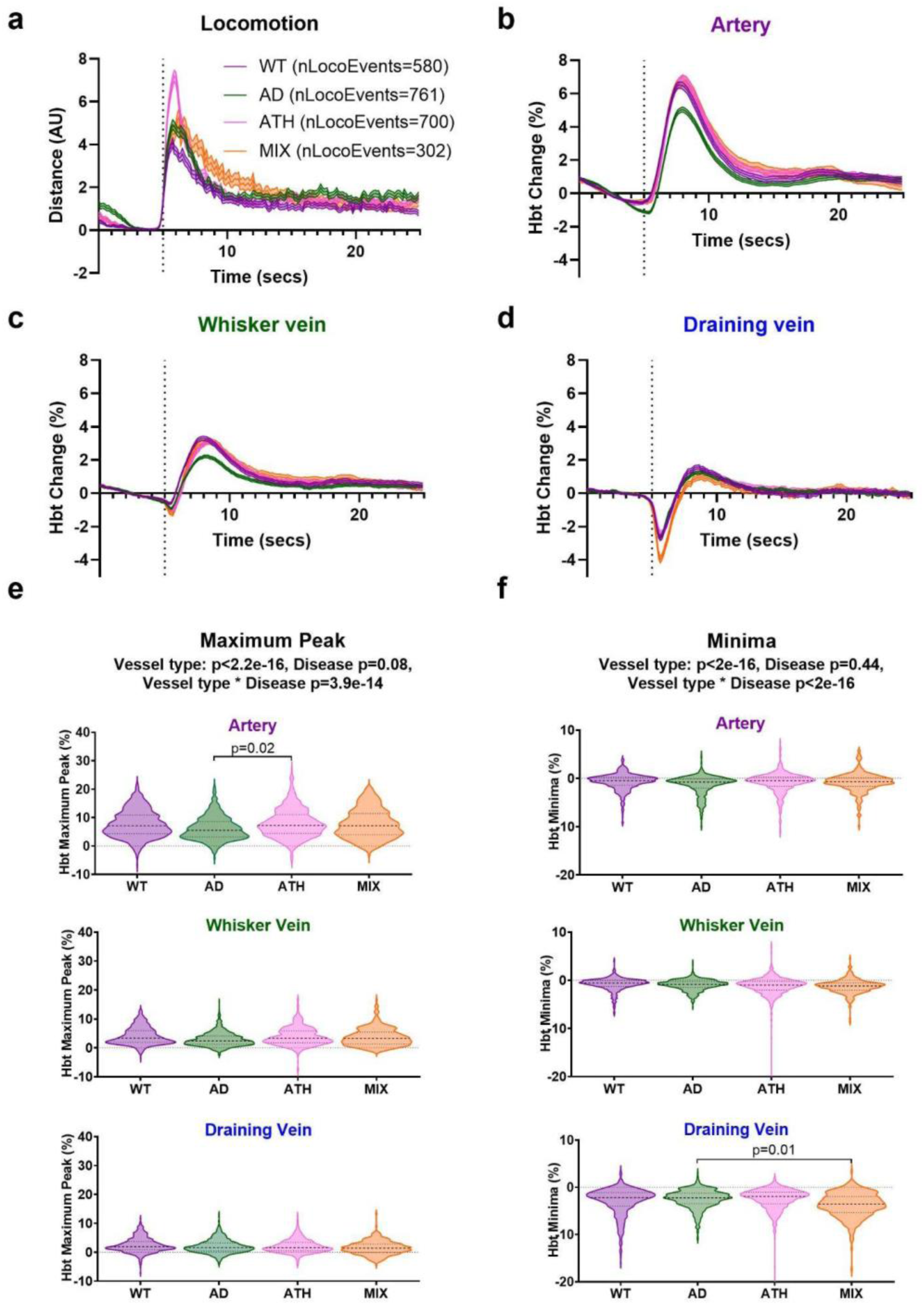
The locomotion-induced draining vein HbT decrease is enhanced in mixed mice. **A.** Locomotion traces were averaged across 4 genotypes (wildtype (WT, purple, nLocoEvents=580), APP/PS1 (AD, green, nLocoEvents=761), Atherosclerosis (ATH, pink, nLocoEvents=700), and APP/PS1 x Atherosclerosis (MIX, orange, nLocoEvents=302)). There were no significant differences in locomotion between genotypes (Vessel type p=1.00, Genotype p=0.33, Vessel type * Genotype p=1.00; assessed using area under the curve from individual locomotion events), so all locomotion events were included in subsequent analyses. Corresponding HbT traces were visualized across genotypes for the **B.** artery, **C.** whisker vein, and **D.** draining vein. Again, significant differences in the size of the HbT **E.** maximum and **F.** minimas were found across vessel types, with the artery showing the largest increase in HbT and draining vein the largest decrease. There was no significant impact of disease on the size (or timing) of these responses, however there was an interaction between vessel type and genotype as APP/PS1 (AD) mice showed a smaller maximum peak in the artery (AD artery (M: 6.21, SD: 4.17) – ATH artery (M: 8.03, SD: 5.02), p=0.02), and smaller minima in the draining vein (AD dv (M: -2.48, SD: 2.03) – MIX dv (M: -3.74, SD: 5.10), p=0.01) HbT responses. P-values are taken from linear mixed-effects models with vessel type and genotype inputted as fixed-effect factors, and animal ID as the random effect (see Statistics Report Table SR7). Shaded error bars represent mean +/- SEM. Horizontal lines on violin plots show median and interquartile range. Source data are provided as a Source Data file.

## 3 Results

### 3.1 Large cerebral draining veins display an initial, fast, early decrease in HbT at the onset of locomotion

When representative spatial maps were generated of the changes in HbT across the cortex 1s prior to and up to 5s after spontaneous locomotion onset (see Figure 1d), a decrease in the large draining vein was observed (blue within the large draining vein), closely followed by an increase in HbT within the whisker barrel region (shown in red, increasing in intensity from 2s post-spontaneous locomotion). Because previous work indicates that locomotion induces a constriction in the dural vessels which is likely a “space-conserving” effect^25^, we investigated whether the fast, initial decrease observed in the HbT of the sagittal draining veins was also driven by a constriction. As these recordings were conducted using 2D-optical imaging spectroscopy, rather than 2-photon microscopy, the spatial resolution was limited, meaning we created an average spatial map for the 2s following the onset of detected locomotion events from one representative imaging session (Figure 1e), and the diameter was taken at the largest in-focus point along the draining vein by calculating the full width half maximum of the pixel intensity along a perpendicular line (Figure 1f).

When we characterized the response of the draining vein to locomotion across all our imaging sessions by looking at the average HbT time series across all detected locomotion events (N=2343, Figure 2a), we confirmed the fast, initial decrease in HbT. This decrease was not shown within the whisker artery or vein, where only an increase in HbT was observed (Figure 2b). There was a significant impact of vessel type on the magnitude and timing of the HbT increase, with the whisker artery showing the largest increase (Figure 2c; Supplementary Table SR1; art vs wv p<0.0001, art vs dv p<0.001), as well as reaching its maximum peak in the fastest time (Figure 2d; SR1; art vs wv p=0.08, art vs dv p=0.01) compared to both the whisker vein and draining vein. The minima (size of the largest decrease in the 5s after locomotion onset) was also impacted by vessel type (Figure 2e; SR1) as the draining vein showed the largest decrease in HbT (dv vs art p<0.0001, dv vs wv p<0.0001), an early response as the minimum HbT peak in this vessel occurred significantly faster than the whisker artery maximum HbT peak (Figure 2f; SR1; p<1.2e-12).

### 3.2 The magnitude of the early HbT decrease in the draining vein is not impacted by the size of the locomotion event, unlike the regional HbT increase

The size of the HbT increase (measured by the maximum peak) was influenced by the magnitude of the locomotion event (measured by the area under the curve for the 5s following locomotion onset) across all vessel types (whisker artery, whisker vein, draining vein), as significant linear correlations were observed (Figure 3a). However, the fast, remote initial HbT decrease in the draining vein (measured by the minima) was not impacted by the size of the locomotion event (Figure 3b; SR2; p=0.98). When the locomotion events were ranked in ascending order, and locomotion and hemodynamic traces from the “bottom 10%” and “top 10%” of locomotion events were extracted (Figure 3c-e), whilst there was a significant impact of locomotion on the HbT responses (SR2; p<2.2e-16), there was also an interaction between vessel type and locomotion, as there was no significant difference in the size of the minima for HbT draining vein responses corresponding to the bottom and top 10% of locomotion events from the Tukey post-hoc comparisons accompanying the linear mixed model (Figure 3f; SR3; p=0.93).

### 3.3 Whilst draining and meningeal veins both show a fast HbT decrease following locomotion onset, the meningeal vein does not show the subsequent HbT increase

For a subset of the imaging sessions (57/118 sessions from 21/40 animals) the meningeal vein was also visible within the cranial window, allowing hemodynamic responses associated with locomotion to be extracted and compared to draining vein responses (Figure 4a,b). The draining vein responses from the same 57 imaging sessions and 1224 locomotion trials (Figure 4a) were compared for the subsequent analyses. There was no difference in the relationship between the HbT minima and the size of the locomotion event between the draining vein and the meningeal vein (Figure 4c), as neither vessel type showed a correlation between these metrics. There was a significant difference in the intercept between vessel types however, as the meningeal vessel showed a larger HbT minima overall (Figure 4c). We further explored this difference in the size of the locomotion-induced hemodynamic response between the meningeal and draining vein, showing that the magnitude of the minima was significantly larger for the meningeal vessel (Figure 4d; SR5; p<0.0001) and occurred relatively more quickly when a normalized speed metric was calculated by dividing the absolute size of the HbT minima by the time to reach this value (Figure 4e; SR5; p<0.0001). Interestingly, the HbT responses also differed with respect to the post-HbT minimum decrease response, as the meningeal vessel did not exceed the baseline (pre-locomotion onset) value, whereas the draining vein showed a subsequent increase in HbT (assessed by area under the curve for the 5s following locomotion onset) (Figure 4f; SR5; p<0.0001).

### 3.4 HbT decrease in the draining vein may be affected by disease state

Where the previous figures collapsed the data across disease groups to allow for systematic investigations into the draining vein phenomenon, we subsequently separated the dataset across these groups to investigate whether disease impacted the locomotion-induced HbT responses in the whisker artery, whisker vein and draining vein. 2D-OIS and locomotion recordings were made across 4 different groups of 9-12 month old mice: wild-type (N=580 locomotion events from 12 mice), Alzheimer’s disease (APP/PS1; N=761 locomotion events from 12 mice), atherosclerosis (rAAV8-mPCSK9-D377Y; N=700 locomotion events from 10 mice), and a mixed AD/atherosclerosis model (N=302 locomotion events from 6 mice). The locomotion events and corresponding HbT responses from the whisker artery, whisker vein, and draining vein were compared across disease groups (Figure *5*a-d). We observed no difference in the size of the locomotion events across groups (SR7, not displayed in figure; LMM: vessel type p=1.0, disease group p=0.33, vessel type * disease group p=1.0). However when comparing the HbT maximum peak value (Figure 5e; SR7), whilst we showed no overall impact of disease (p=0.08), we did observe an effect of vessel type (p<2.2e-16) and a vessel type * disease group interaction (p=3.9e-14), likely because the AD mice showed a smaller HbT increase than the atherosclerosis group for the HbT maximum peak in the whisker artery vessel (p=0.02). When we compared the HbT minima (Figure 5f; SR7), whilst we again found no overall impact of disease on the magnitude of this response (p=0.44), we saw an overall effect of vessel type on the size of the minimum HbT decrease (p<2.2e-16), and a vessel type * disease group interaction (p<2.2e-16) which post-hoc comparisons revealed was due to a smaller response in AD mice compared to the mixed model in the draining vein (p=0.01).

## 4 Discussion

### 4.1 The “B of the Bang” for neurovascular coupling

In this study we collected hemodynamic responses to spontaneous locomotion in awake, head-fixed mice whilst they walked on a spherical treadmill. Surprisingly, we observed that large cerebral draining veins had a rapid and early decrease in HbT at the onset of locomotion. This observation occurred prior to increases in HbT within the pial artery and vein in the whisker barrel region and, unlike these HbT increases, was not impacted by the magnitude of the locomotion event. Given the rapid nature of this HbT decrease, which precedes the eventual HbT influx induced by locomotion, we propose it as the “B of the Bang” for neurovascular coupling (a phrase coined by British sprinter Linford Christie, when he emphasized he does not start his races merely at the “bang” of the pistol)^45^. The HbT decrease observed in the draining vein was also different to that seen in meningeal vessels (as previously reported by Gao et al., 2016^25^), as the meningeal vessels did not show the subsequent increase in HbT above baseline levels associated with the impact of locomotion. We also assessed whether this initial decrease in HbT within the draining vein was affected by pathological conditions, observing that disease group and vessel type interacted for both the HbT maximum peak and minima, as AD mice showed a smaller HbT increase in the whisker artery versus atherosclerosis mice, and a smaller HbT minima in the draining vein versus the mixed AD/atherosclerosis mice.

### 4.2 Comparison to previous studies

Whilst other groups investigating the impact of locomotion on pial and dural vessels have reported the dural constriction at the onset of locomotion^25^, until now no other studies have reported the fast, initial HbT decrease in the large draining veins of the sagittal sinus. Previous work investigating the locomotion-induced meningeal constriction showed that the response in these vessels was related to changes in intracranial pressure, and was distinct from pial vessel responses^25^. In a follow-up study from the same group, a widefield imaging approach (similar to 2D-OIS) was used to assess the longitudinal effects of locomotion and respiration on cerebral oxygenation^46^. Whilst the authors do not report an initial decrease in blood volume in the draining veins at the onset of locomotion, representative spatial images shown in their Figure 2f indicate the draining vein (bottom of image) having a response very similar to those observed in our study.

Additionally, Zhang et al^46^ developed a new method of fast kilohertz two-photon imaging in awake mice. As part of their methodological approach they investigated the diameter of the large pial veins close to the sagittal sinus (i.e. the draining veins). They reported sharp constrictions in vein diameter in the awake animal at multiple time points but did not correlate this with behaviors such as locomotion. Taken together, these results suggest that, in response to locomotion, prior to a large increase in bilateral perfusion, large pial draining veins reduce their size. Further research using bilateral cranial windows and simultaneous measurements of neuronal activity, potentially using GCAMP imaging technologies will help understand this response further with respect to brain metabolism.

Studying the large draining and meningeal veins has important implications for enhancing understanding of the brain’s glymphatic waste drainage system, as it is postulated that waste may be cleared along the venous circulation^47,48^ to the downstream lymphatic network (associated with the large sagittal vessels exiting to the meninges)^49–51^. Although this clearance process is not fully understood and controversies surrounding the existing mechanisms still exist, the oscillation of blood vessel diameter is thought to be important, as an interruption of vascular oscillations results in reduced clearance^52^, and may lead to an increase in Aβ accumulation^53^.

Hence investigations of the dilation capacity and blood volume responses across the relevant vascular segments may enhance our understanding of the mechanisms relevant for glymphatic drainage in aging and neurodegeneration. In this study we show an effect of disease in the draining vein locomotion-induced haemodynamic response, as mixing atherosclerosis and AD resulted in a larger HbT decrease versus in the AD only model, which could have important implications for the impact of co-morbid disease mechanisms on glymphatic drainage systems.

### 4.3 Limitations

Whilst we hypothesized that the early decrease in HbT within the large draining veins of the brain potentially occurred due to making more ‘space’ within the brain for the subsequent large increases in HbT induced by spontaneous locomotion, as we did not measure intracranial pressure during these experiments we cannot make any causal inferences about pressure changes within the brain when these early responses occur. However, Gao and colleagues (2016)^25^ did conduct extensive experiments assessing ICP changes in awake, behaving mice. They reported that ICP increases were observed during locomotion, these occurred rapidly and returned to baseline values once locomotion ceased. They further reported that when fitting the data to a linear convolution model the ICP increases occurred just prior to locomotion. Therefore, it could be suggested that the decrease in HbT within the large draining veins may be explained by increases in ICP compressing these large veins to make space for the subsequent large increase in HbT that follows locomotion. As we used 2D-Optical imaging spectroscopy, we were unable to explicitly measure whether there was a constriction across all the draining veins recorded (due to limited spatial resolution). Therefore, it would be informative to investigate the above findings using 2-photon microscopy, and in future follow-up studies we plan to explicitly measure vessel dynamics and investigate absolute blood flow changes within these vessels.

We also had no measure of neural activity within the study, meaning we could not capture brain metabolism changes in response to locomotion comprehensively. It would be beneficial to establish how neural activity is affected by spontaneous locomotion and the relationship this may have with the draining vein. Other studies have investigated how neural activity is impacted by spontaneous locomotion^20,24^. However, it would be especially interesting to focus on neuronal responses close to the draining vein. For instance, future work to conduct similar experiments using GCaMP mice^51^ in order to investigate neural activity and hemodynamics concurrently in awake mice may give a deeper insight on neural activity corresponding to hemodynamic responses within the draining veins of the brain.

Furthermore, beyond just locomotion, it has been suggested that it is important to track the range of behaviors of the awake mouse such as arousal state, fidgeting, whisking, pupil diameter, or breathing rate, which have all been implicated as important modulators of cerebral hemodynamic activity^22,46,54,55^, and in the case of breathing rate may impact oxygenation status independently of underlying neural activity^46^.

It has also been suggested that very early vascular responses may represent artifacts, potentially as a result of the startle response of the animals^56–58^. The animals used in this study underwent extensive habituation to the experimental set-up prior to recording, and the spontaneous locomotion trials were created from individual walking events during a trial in which there were no whisker stimulations. It would be odd to suggest that the animal walking would cause itself to startle. Locomotion is not a reflex response and in order for the mouse to move, the animal has to initiate this response^59^, therefore making it very unlikely that these findings are the effects of the startle response.

### 4.4 Conclusions

Our paper demonstrates the importance of monitoring locomotion during awake hemodynamic imaging, as we have shown that spontaneous locomotion onset impacts hemodynamic responses differentially across the vascular network. Understanding where and how different vascular functions are supported across the vascular tree is vital to understanding how the different sections may be impacted, or targeted, during disease or aging. We suggest that this HbT decrease observed in the draining veins may be an important ‘space-saving’ mechanism involved in NVC and may allow for the large increase in HbT that follows locomotion, and that this mechanism is enhanced in mixed model atherosclerosis/AD mice (compared to AD only). This finding relating to an increased HbT decrease in the draining vein of the mixed disease model could be particularly relevant for the impact of disease on glymphatic clearance pathways. Further studies are warranted using 2-photon microscopy to confirm whether the early reduction in HbT within the draining vein is the result of a constriction of large draining veins at the onset of locomotion. This research shows that large cerebral veins may be important components of the neurovascular unit, rather than passive bi-standers or balloons waiting to be inflated.

## 6 Code, Data, and Materials Availability

- The data presented in this article are publicly available on ORDA at [DOI link to be included before publication]. The individual hemodynamic and locomotion traces with corresponding labels (vessel group, disease group, imaging session, locomotion event number) are available as MATLAB files, and the averaged traces presented in the Figures and time series parameters used for the statistical calculations as a .csv file.
- The custom code used for data analysis is available as a GitHub repository, and has been published via Zenodo (DOI: 10.5281/zenodo.14223430).
- The outputs of all statistical analyses are reported in statistical tables in Appendix A.

## Acknowledgements

We thank Rachel Sandy at the University of Sheffield for her technical expertise with IV injections as well as the entire BSU team for their husbandry expertise. CH was funded by a Sir Henry Dale Fellowship jointly funded by the Wellcome Trust and the Royal Society. This research was funded in whole, or in part, by the Wellcome Trust [Grant number 105586/Z/14/Z]. The research was also funded by a Batelle-Jeff Wadsworth PhD Studentship (BE) and a UKRI MRC grant [Grant number MR/X003418/1]. For the purpose of open access, the author has applied a Creative Commons Attribution (CC BY) license to any Author Accepted Manuscript version arising. Some figures were Created in https://BioRender.com.

## Author Biographies

**Beth Eyre** is a research fellow at Massachusetts General Hospital/Harvard Medical School. She received her BSc degree (Psychology, *International*) from the University of Leeds in 2017, her MSc in Cognitive Neuroscience and human Neuroimaging from the University of Sheffield in 2020 and also her PhD in 2023. She is an author on 4 journal papers and her current research interests include cerebral amyloid angiopathy, brain clearance and neurovascular coupling.

**Kira Shaw** is a postdoctoral research associate at the University of Sheffield. She received her BS degree in Psychology from the University of Leeds in 2011, her MS Psychology degree from the University of Manchester in 2012, and her PhD in Neuroscience from the University of Sheffield in 2017. She is an author on 10 journal papers, and her current research interests include studying neurovascular coupling and the impact perturbations in energy supply may place on brain function.

**Jason Berwick** is a Reader in Neurophysiology and Neuroimaging at the University of Sheffield who specializes in the use of in vivo multimodal brain imaging and electrophysiological methodologies to understand the mechanisms of neurovascular coupling in health and disease. He has published over 60 research papers on the topic of NVC.

**Clare Howarth** is a Senior Lecturer at the University of Sheffield. She received her MSci degree in Physics with a year in Europe from Imperial College London in 2003, and her PhD in Neuroscience from University College London in 2008. She is an author on 29 papers, and her current research interests focus on neurovascular coupling in health and disease.

Biographies and photographs for the other authors are not available.

