## Supplementary data for "Voluntary locomotion induces an early and remote hemodynamic decrease in the large cerebral veins"

**Appendix A: Statistical Reports**

**Statistics Reports (SR1-SR7):** The following tables report the mean, standard deviation, type of statistical test, test statistics, degrees of freedom and p-value for each statistical test reported in the main manuscript.

**SR1: Figure 2: Spontaneous locomotion induces a fast, remote decrease in the large cerebral draining vein that precedes the regional HbT influx.** The three level comparisons (vessel type: artery, whisker vein, draining vein) for 2c-e were subject to a linear mixed model analysis (lmer package RStudio), with vessel type entered as the fixed factor and animal ID as the random factor (to account for variations between groups being driven by a single outlier animal). To further investigate where the specific significant differences were pairwise comparisons (with correction for multiple comparisons) were conducted using the Tukey method (emmeans function RStudio). The two group comparison for 2f was first subject to a Shapiro-Wilk test (shapiro.test function RStudio) to assess normality, and due to a significant result indicating data was not normally distributed, compared using the non-parametric Wilcoxon-Signed ranks test for paired data (wilcox.test function in RStudio, paired = true).

| **Figure** | **Mean** | **SD** | **Test** | **Test Statistic** | **Mean Square** | **Degrees of freedom** | **P Value** |
|---|---|---|---|---|---|---|---|
| 2c (Hbt max peak) | Art: 7.33<br>WV: 3.61<br>DV: 2.06 | Art: 4.77<br>WV: 2.89<br>DV: 2.40 | Linear mixed model (fixed factor: vessel type) | F = 1668.1 | 16991 | 2 | p<2.2e-16<br><br>**Pairwise**<br>art-dv: p<.0001<br>art-wv: p<.0001<br>dv-wv: p<.0001 |
| 2d (Hbt time to max peak) | Art: 3.13<br>WV: 3.19<br>DV: 3.22 | Art: 0.96<br>WV: 1.03<br>DV: 1.27 | Linear mixed model (fixed factor: vessel type) | F=4.51 | 5.06 | 2 | p=0.011<br><br>**Pairwise**<br>art-dv: p=0.011<br>art-wv: p=0.082<br>dv-wv: p=0.726 |
| 2e (Hbt minimum peak) | Art: -0.97<br>WV: -1.06<br>DV: -2.77 | Art: 2.01<br>WV: 1.73<br>DV: 2.56 | Linear mixed model (fixed factor: vessel type) | F=582.64 | 2414.1 | 2 | p<2.2e-16<br><br>**Pairwise**<br>art-dv: p<.0001<br>art-wv: p=0.291<br>dv-wv: p<.0001 |
| 2f (compare peak timing (max for art, min for dv) | Art: 3.13 | Art: 0.96 | Paired Wilcoxon-Signed Rank | W=269 | | 1 | p<2.2e-16 |

| | | |
|---|---|---|
| | DV: 1.04 | DV: 0.94 |

**SR2: Figure 3: The amount of locomotion modulates the size of the HbT increase, but not the initial HbT decrease in the draining vein.** For correlation analyses in 3a-b, p-values are taken from a Pearson's correlation (cor.test package RStudio) which tested for a linear relationship between the size of the locomotion event (AUC for 5s after onset) and the size of the HbT maximum or minimum peak (calculated from the maximum or minimum value reached in 5s after locomotion onset).

| **Figure** | **Test** | **Test Statistic** | **95% CI** | **Degrees of freedom** | **P Value** |
|---|---|---|---|---|---|
| 3a (Art maxpk) | Pearson's Correlation | t=14.48 r=0.28 | 0.249 - 0.323 | 2341 | p<2.2e-16 |
| 3a (WV maxpk) | Pearson's Correlation | t=10.98 r=0.22 | 0.182-0.259 | 2341 | p<2.2e-16 |
| 3a (DV maxpk) | Pearson's Correlation | t=7.49 r=0.15 | 0.113-0.192 | 2341 | p=9.6e-14 |
| 3b (Art minpk) | Pearson's Correlation | t=-2.64 r=-0.05 | -0.095–0.014 | 2341 | p=0.0083 |
| 3b (WV minpk) | Pearson's Correlation | t=-2.23 r=-0.05 | -0.086–0.00056 | 2341 | p=0.0257 |
| 3b (DV minpk) | Pearson's Correlation | t=-0.02 r=-0.00039 | -0.041-0.040 | 2341 | p=0.9849 |

**SR3: Figure 3: The amount of locomotion modulates the size of the HbT increase, but not the initial Hbt decrease in the draining vein.** A linear mixed model was conducted for 3f which tested for a significant impact of vessel type (3 levels: artery, whisker vein, draining vein), locomotion group (2 levels: bottom or top 10% locomotion trials) or a vessel type * locomotion group interaction on HbT peak responses (maximum peak for artery and wv, minimum peak for dv). Vessel and locomotion groups were inputted as fixed factors, and animal ID as the random factor using the lmer package in RStudio.

| **Figure** | **Mean** | **SD** | **Test** | **Test Statistic** | **Mean Square** | **Degrees of freedom** | **P Value** |
|---|---|---|---|---|---|---|---|
| 3f (peak) | Art maxpk bottom | Art maxpk bottom 10%: 2.77 | Linear mixed model | Vessel type F=1478.29, | Vessel type 10084.1, Locomotion | Vessel type 2, Locomotion | Vessel type p<2.2e-16, Locomotion |

| | | | | | | | |
|---|---|---|---|---|---|---|---|
| | 10%: 4.61 Art maxpk top 10%: 9.49 WV maxpk bottom 10%: 2.06 WV maxpk top 10%: 4.61 DV minpk bottom 10%: -1.79 DV minpk top 10%: -2.56 | Art maxpk top 10%: 5.11 WV maxpk bottom 10%: 1.52 WV maxpk top 10%: 3.19 DV minpk bottom 10%: 1.79 DV minpk top 10%: 2.21 | (fixed factors: vessel type, locomotion group) | Locomotion group F=134.19, Vessel type * Locomotion group F=138.52 | group 915.3, Vessel type * Locomotion group 944.9 | group 1, Vessel type * Locomotion group 2 | group p<2.2e-16, Vessel type * Locomotion group p<2.2e-16 |
| Pairwise comparisons: | bottom 10% art - top 10% art p<.0001, bottom 10% art - bottom 10% dv p<.0001, bottom 10% art - top 10% dv p<.0001, bottom 10% art - bottom 10% wv p<.0001, **bottom 10% art - top 10% wv p=0.6273**, top 10% art - bottom 10% dv p<.0001, top 10% art - top 10% dv p<.0001, top 10% art - bottom 10% wv p<.0001, top 10% art - top 10% wv p<.0001, **bottom 10% dv - top 10% dv p=0.9254**, bottom 10% dv - bottom 10% wv p<.0001, bottom 10% dv - top 10% wv p<.0001, top 10% dv - bottom 10% wv p<.0001, top 10% dv - top 10% wv p<.0001, bottom 10% wv - top 10% wv p<.0001 **Non-significant comparisons denoted in bold font** | | | | | | |

**SR4: Figure 3: The amount of locomotion modulates the size of the HbT increase, but not the initial HbT decrease in the draining vein.** As well as conducting two-group comparisons using the Tukey post-hoc (emmeans package in RStudio) associated with the linear mixed model, we also analysed the maximum peak (for whisker artery and whisker vein) independently of the minimum peak in the draining vein. For the maximum peak we conducted a non-parametric two-way ANOVA (ggpubr package in RStudio) due to the data not being normally distributed (Shapiro-Wilks test: art- W=0.91, p=9.02e-7; wv- W=0.85, p=1.31e-9), and for the minimum peak in the draining vein we conducted a Wilcoxon rank sum test due to the data not being normally distributed (Shapiro-Wilks test: dv- W=0.88, p=4.92e-8).

| Figure | Mean | SD | Test | Test Statistic | Mean Square | Degrees of freedom | P Value |
|---|---|---|---|---|---|---|---|
| 3f (maximum peaks) | See SR3 | See SR3 | Non-parametric two-way ANOVA | Vessel Type F=89.44, Locomotion group F=142.97, Vessel type * Locomotion group F=11.47 | Vessel Type 744.97, Locomotion group 1190.86, Vessel type * Locomotion group 95.53 | Vessel Type 1, Locomotion group 1, Vessel type * Locomotion group 1 | Vessel Type p<2.2e-16, Locomotion group p<2.2e-16, Vessel type * Locomotion |

| | | | | | | | |
|---|---|---|---|---|---|---|---|
| | | | | | | | group p=0.00083 |
| Pairwise comparisons: | Art bottom – wv bottom p<0.0001, art bottom – art top p<0.0001, art bottom – wv top p=0.467, wv bottom – art top p<0.0001, wv bottom – wv top p<0.0001, art top – wv top p<0.0001 | | | | | | |
| 3f (dv minimum peak) | See SR3 | See SR3 | Paired Wilcoxon-Signed Rank | W=2034 | | 1 | p=0.024 |

**SR5: Figure 4: The meningeal vein shows a larger HbT decrease following locomotion onset, which unlike for the draining vein does not subsequently increase above baseline.** Two-group comparisons were conducted to test for differences in HbT dynamics between draining and meningeal vessels. The assumptions of the unpaired t-test were assessed in GraphPad Prism using a Kolmogorov-Smirnov test of normality, and F test to compare the variances. For each figure plot (4c-f) the assumptions were violated, and the Mann-Whitney non-parametric alternative test was used.

| **Figure** | **Mean** | **SD** | **Test** | **Test Statistic** | **Sum of Ranks** | **Degrees of freedom** | **P Value** |
|---|---|---|---|---|---|---|---|
| 4d HbT Minima | dv: -2.358 mn: -3.971 | dv: 1.952 mn: 3.063 | Mann-Whitney test | U= 502017 | dv: 1745859 mn: 1251717 | 1222 | p<0.0001 |
| 4e HbT Normalised speed metric | dv: 0.9924 mn: 4.544 | dv: 0.912 mn: 4.656 | Mann-Whitney test | U= 216212 | dv: 965912 mn: 2031664 | 1222 | p<0.0001 |
| 4f HbT Area Under Curve | dv:0.7059 mn: -6.105 | dv: 8.44 mn: 12.73 | Mann-Whitney test | U= 471610 | dv: 1776266 mn: 1221310 | 1222 | p<0.0001 |

**SR6: Figure 4: The meningeal vein shows a larger HbT decrease following locomotion onset, which unlike for the draining vein does not subsequently increase above baseline.** Due to the significant difference in the amount of locomotion between draining vein and meningeal groups, the impact of locomotion on the Hbt minima was assessed using a linear regression. Neither the draining vein nor the meningeal vessel showed a significant correlation between locomotion and the HbT minimum (see is the slope significantly non-zero). Furthermore, there was no significant difference between the vessel groups on the relationship between locomotion and the HbT minima (see are the slopes equal). The intercepts were significantly different between groups as the minimum peak negative values were larger in the meningeal group.

| **Figure** | **Test** | **Is the slope significantly non-zero?** | **Are the slopes equal?** | **Are the intercepts equal?** |
|---|---|---|---|---|

| | | | | |
|---|---|---|---|---|
| 4c Locomotion / HbT Minima Scatter | Linear Regression | dv: F(1,1222)=0.3038, p=0.5816<br>mn: F(1,1222)=0.5123, p=0.474 | F(1,2444)=0.0944, p=0.7587 | F(1,2445)=241.3, p<0.0001 |

**SR7: Figure 5: The locomotion-induced draining vein HbT decrease is enhanced in mixed mice.** A linear mixed model was conducted for 5e-f which tested for a significant impact of vessel type (3 levels: artery, whisker vein, draining vein), genotype (4 levels: group 1, group 2, group 3, group 4) or a vessel type * genotype interaction on HbT peak responses (HbT maximum peak for e, HbT minimum peak for f). Vessel and genotype groups were inputted as fixed factors, and animal ID as the random factor using the lmer package in RStudio.

| **Figure** | **Mean** | **SD** | **Test** | **Test Statistic** | **Mean Square** | **Degrees of freedom** | **P Value** |
|---|---|---|---|---|---|---|---|
| 5e (HbT max peak) | Art: AD: 6.21 WT: 7.75 ATH: 8.00 MIX: 7.81 WV: AD: 2.92 WT: 4.11 ATH: 3.88 MIX: 3.75 DV: AD: 2.00 WT: 2.43 ATH: 2.08 MIX: 1.49 | Art: AD: 4.17 WT: 4.76 ATH: 5.02 MIX: 5.10 WV: AD: 2.38 WT: 2.97 ATH: 3.11 MIX: 3.07 DV: AD: 2.34 WT: 2.53 ATH: 2.34 MIX: 2.32 | Linear mixed model (fixed factors: vessel type, genotype) | Vessel type F=1569.7, Genotype F=2.372, Vessel type * Genotype F=12.556 | Vessel type 15832.3, Genotype 23.9, Vessel type * Genotype 126.6 | Vessel type 2, Genotype 3, Vessel type * Genotype 6 | Vessel type p<2.2e-16, Genotype p=0.086, Vessel type * Genotype p=3.9e-14 |
| Pairwise comparisons: | **Due to high number of comparisons, only non-significant results reported:** art AD - art WT p=0.128, art AD - wv WT p=0.103, art AD - wv ATH p=0.190, art AD - art MIX p=0.208, art AD - wv MIX p=0.288, dv AD - dv WT p=0.989, dv AD - dv ATH p=0.999, dv AD - wv MIX p=0.129, wv AD - wv WT p=0.435, wv AD - wv ATH p=0.473, wv AD - dv MIX p=0.986, wv AD - wv MIX p=0.856, art WT - art ATH p=0.999, art WT - art MIX p=1.00, dv WT - dv ATH p=1.00, dv WT - wv ATH p=0.266, dv WT - dv MIX p=0.998, dv WT - wv MIX p=0.702, wv WT - dv ATH p=0.208, wv WT - wv ATH p=1.00, wv WT - wv MIX p=1.00, art ATH - art MIX p=1.00, dv ATH - dv MIX p=0.999, dv ATH - wv MIX p=0.643, wv ATH - wv MIX p=1.00 | | | | | | |

| | | | | | | | |
|---|---|---|---|---|---|---|---|
| | **Note that for interpreting vessel type * genotype interactions attention was focused on within vessel type comparisons that were significantly different between genotype (i.e. for artery max peak group 1 and group 3 differed p=0.02).** | | | | | | |
| 5f (HbT min) | Art:<br>AD:-1.37<br>WT:-0.71<br>ATH:-0.82<br>MIX:-0.82<br>WV:<br>AD:-0.98<br>WT:-0.81<br>ATH:-1.26<br>MIX:-1.29<br>DV:<br>AD:-2.48<br>WT:-2.92<br>ATH:-2.54<br>MIX:-3.74 | Art:<br>AD: 2.03<br>WT: 1.73<br>ATH: 2.14<br>MIX: 2.01<br>WV:<br>AD: 1.13<br>WT: 1.32<br>ATH: 2.49<br>MIX: 1.45<br>DV:<br>AD: 1.98<br>WT: 2.84<br>ATH: 2.72<br>MIX: 2.63 | Linear mixed model (fixed factors: vessel type, genotype) | Vessel type F=626.83, Genotype F=0.927, Vessel type * Genotype F=22.684 | Vessel type 2549.11, Genotype 3.77, Vessel type * Genotype 92.25 | Vessel type 2, Genotype 3, Vessel type * Genotype 6 | Vessel type p<2e-16, Genotype p=0.438, Vessel type * Genotype p<2e-16 |
| Pairwise comparisons: | **Due to high number of comparisons, only non-significant results reported:** art AD - art WT p=0.838, art AD - wv WT p=0.949, art AD - art ATH p=0.999, art AD - wv ATH p=1.00, art AD - art MIX p=0.999, art AD - wv MIX p=1.00, dv AD - dv WT p=0.757, dv AD - dv ATH p=0.999, dv AD - wv ATH p=0.146, dv AD - wv MIX p=0.442, wv AD - art WT p=1.00, wv AD - wv WT p=1.00, wv AD - art ATH p=1.00, wv AD - wv ATH p=0.941, wv AD - art MIX p=1.00, wv AD - wv MIX p=0.971, art WT - wv WT p=0.999, art WT - art ATH p=0.999, art WT - wv ATH p=0.658, art WT - art MIX p=1.00, art WT - wv MIX p=0.808, dv WT - dv ATH p=0.998, dv WT - dv MIX p=0.555, wv WT - art ATH p=1.00, wv WT - wv ATH p=0.835, wv WT - art MIX p=1.00, wv WT - wv MIX p=0.916, art ATH - wv MIX p=0.993, dv ATH - dv MIX p=0.135, dv ATH - wv MIX p=0.119, wv ATH - art MIX p=0.997, wv ATH - wv MIX p=1.00, art MIX - wv MIX p=0.150. **Note that for interpreting vessel type * genotype interactions attention was focused on within vessel type comparisons that were significantly different between genotype (i.e. for dv min peak group 1 and group 4 differed p=0.01).** | | | | | | |
